# Dynamics of absolute and relative disparity processing in human visual cortex

**DOI:** 10.1101/2021.04.15.440053

**Authors:** Milena Kaestner, Marissa L. Evans, Yulan D. Chen, Anthony M. Norcia

## Abstract

Cortical processing of binocular disparity is believed to begin in V1 where cells are sensitive to absolute disparity, followed by the extraction of relative disparity in higher visual areas. While much is known about the cortical distribution and spatial tuning of disparity-selective neurons, the relationship between their spatial and temporal properties is less well understood. Here, we use steady-state Visual Evoked Potentials and dynamic random dot stereograms to characterize the temporal dynamics of spatial mechanisms in human visual cortex that are primarily sensitive to either absolute or relative disparity. Stereograms alternated between disparate and non-disparate states at 2 Hz. By varying the spatial frequency content of the disparate fields from a planar surface to corrugated ones, we biased responses towards absolute vs. relative disparities. Reliable Components Analysis was used to derive two dominant sources from the 128 channel EEG records. The first component (RC1) was maximal over the occipital pole while the second component (RC2) was maximal over right lateral occipital electrodes. In RC1, first harmonic responses were sustained, tuned for corrugation frequency, and sensitive to the presence of disparity references, consistent with prior psychophysical sensitivity measurements. By contrast, the second harmonic, associated with transient processing, was not spatially tuned and was indifferent to references, consistent with it being generated by an absolute disparity mechanism. In RC2, the sustained response component showed similar tuning and sensitivity to references. However, sensitivity for absolute disparity dropped off, and transient signals were mainly driven by the lowest corrugation frequencies.

**SIGNIFICANCE STATEMENT:** Sustained and transient mechanisms have been demonstrated across sensory systems and may reflect a common coding strategy. This strategy may be useful for the analysis of the form and duration of complex events in the case of sustained signals, and the onset and location of events in the case of transient signals. Here, we provide direct neural correlates of sustained and transient disparity mechanisms in human visual cortex. Early visual cortex is sensitive to both relative and absolute disparity, with the former being processed in a sustained fashion and the latter in a more transient fashion. Outside of early visual cortex, sustained relative disparity responses are readily measurable, but transient responses and responses to absolute disparity are not.

## INTRODUCTION

In the perceptual and oculomotor literatures, at least four functional dichotomies have been proposed to underly the percept of depth from disparity. These include processes common to other visual modalities, such as local or global processing, (Julesz, 1971), coarse or fine mechanisms (Wilcox and Allison, 2009), first-order vs. second-order processing (Hess and Wilcox, 1994), and temporally transient or sustained mechanisms (Westheimer and Mitchell, 1969; Mitchell, 1970; Jones, 1980; Edwards et al., 1998). Other stimulus-based dichotomies specific to stereopsis include absolute vs. relative disparity, crossed vs. uncrossed disparities, and horizontal vs. vertical disparities.

It is likely that some of these functional and stimulus-based dichotomies are inter-related, sharing a common set of neural resources. It is therefore desirable to identify a smaller number of component processes that can be mapped onto underlying neural mechanisms. In this study, we aim to identify neural processes underlying the spatial stimulus constructs of absolute and relative disparity and to unify them with the temporal functional constructs of transient and sustained mechanisms.

Absolute and relative disparity are computationally distinct and appear to be processed by different mechanisms. Absolute disparity is the difference in angle subtended on the left and right retina of an object in space and gives an estimate of the depth of that object to the observer. Relative disparity is the comparative depth between two objects in space and arises when there are two or more depth planes present. Perceptually, depth judgements are dominated by relative disparity – observers are able to discriminate smaller changes in disparity in the presence of a reference than in its absence (Westheimer, 1979; McKee et al., 1990; Kumar and Glaser, 1991; Andrews et al., 2001). Stereoacuity, a relative disparity task, improves with increasing exposure duration up to several seconds (Ogle and Weil, 1958; Harwerth and Rawlings, 1977), suggesting that it is subserved by neural mechanisms that are sustained.

Studies of the vergence system have implicated both transient and sustained disparity-tuned processes. Vergence can be initiated by disparate, but not fusible targets that vary in their absolute disparity, but only fusible targets allow for a sustained vergence response. (Westheimer and Mitchell, 1969; Mitchell, 1970; Jones, 1980). While changes in absolute disparity over time provide a strong cue for vergence eye movements, they lead to weak percepts of motion-in-depth (Erkelens and Collewijn, 1985a, b; Regan et al., 1986; Cottereau et al., 2012b).

Behavioural findings imply that neurons tuned for absolute disparity have transient responses, while neurons responsive to relative disparity have sustained responses. To our knowledge, there are no comparative studies of the temporal dynamics of cell responses to absolute vs. relative disparities. One study in V1 of macaque, where cells are sensitive to absolute disparity, has suggested that these responses are sustained (Nienborg et al., 2005).

The terms ‘transient’ and ‘sustained’ have been used in the literature to refer to properties of the stimulus, the resulting visual percept, or the underlying neural mechanisms (Gheorghiu and Erkelens, 2005). Psychophysical studies have necessarily depended on manipulations of the stimulus to infer dynamics. Here, we take a different approach and capitalise on the temporal resolution of EEG to acquire a direct neural readout. Using dynamic random dot stereograms to generate steady-state visual evoked potentials, we isolate neural responses to changing disparity. By varying the availability of disparity references and the corrugation frequency of the target, we vary the availability of absolute vs. relative disparity cues without changing the monocular image content.

Using a spatial filtering approach (Dmochowski et al., 2015), we identify a neural source whose sustained response component is strongly tuned for corrugation frequency, but whose transient component is not. A second source, maximal over right occipital electrodes, shows a similar response pattern in the sustained component whilst absolute disparity sensitivity drops off. We conclude that there is a dominant sustained channel that is strongly tuned for corrugation frequency and thus relative disparity, whilst the transient channel is best driven by absolute disparity.

## METHODS

### Participants

All participants were recruited from the Stanford community, and were screened for normal or corrected-to-normal vision, ocular diseases and neurological conditions. Visual acuity was measured using a LogMAR chart (Precision Vision, Woodstock, IL, USA) and was better than 0.1 in each eye with less than 0.3 acuity difference between the eyes. Stereoacuity was measured with the RANDOT stereoacuity test (Stereo Optical Company, Inc., Chicago, IL, USA) with a pass score of 50 arcsec or better. In the first experiment (reference effects), 22 participants (13 female, 9 male, mean age 31 years) were recruited. Data from one participant were excluded for technical issues during the recording, and data from a second participant were excluded for a low signal-to-noise ratio in the EEG response component indexing low-level luminance changes (dot update response occurring at 20 Hz, see Methods: Visual Display for more detail). Data from the remaining 20 participants were retained for analysis. In the second experiment (corrugation tuning), 30 participants (15 female, 15 male, mean age 34 years) were recruited. Of these, five were excluded from analysis, two due to ocular and other chronic diseases that met the exclusion criteria and two for technical issues arising during the recording. One participant was excluded for a low signal-to-noise ratio in the dot update response. Data from 25 participants were retained for analysis. Informed written and verbal consent was obtained from all participants prior to participation under a protocol approved by the Institutional Review Board of Stanford University.

### Visual Display

Stimuli were displayed on a SeeFront 32” autostereoscopic 3D monitor running at a refresh rate of 60 Hz. The SeeFront display comprises a TFT LCD panel with an integrated lenticular system that interdigitates separate images for the left and right eyes on alternate columns of the 3840 x 3160 native display resolution. In 3D mode, the effective resolution in 1920 x 1080 pixels per eye. Mean luminance was 50 cd/m^2^ as requested by the stimulus generation software after in-house calibration and gamma linearization. The viewing distance was 70 cm which is within the optimal range for the adult average of a 65 mm inter-pupillary distance, as per the manufacturer’s specifications. At this distance, the total field of view in degrees of visual angle was 53.8° x 31.5°. The SeeFront device monitors the participants’ head position via an integrated pupil location tracker and shifts the two eyes’ views to compensate for motion, thus ensuring that the images are projected separately into each eye. Head positioning was checked periodically for each participant by asking them to report on the separate visibility of the nonius lines.

The stimulus comprised dynamic random-dot stereograms (DRDS) whose frames were generated in MATLAB using Psychtoolbox-3 (Brainard, 1997; Pelli, 1997; Kleiner et al., 2007). These frames were presented via a custom Objective C application with no jitter or frame dropping. The general layout of each stimulus condition is illustrated in Figure 1, Panel A. The DRDS were viewed through a circular aperture (28.5° diameter) embedded within a square 39.5° by 39.5° 1/f noise fusion lock that was used to control eye gaze and vergence angle. The fusion lock was at zero disparity (identical images in left and right eyes). A ring of binocularly uncorrelated dots that was 1.2° wide was placed in between the edge of the DRDS and the fusion lock to reduce the availability of relative disparity cues arising from the edge of the zero-disparity fusion lock and the DRDS (Cottereau et al., 2012a). These uncorrelated dots were identical to the stimulus dots in size, density and contrast but their positions did not update, and they were static during each stimulus trial. The edge between the uncorrelated dots and the DRDS was blended with a cosine ramp, and the visible diameter of the DRDS stimulus was 27.3°. To control the eye position of participants, and to aid stable binocular fusion, nonius lines were placed at central fixation where the length of each line was 1° with 0.3° separation between upper and lower lines.

**Figure 1:**
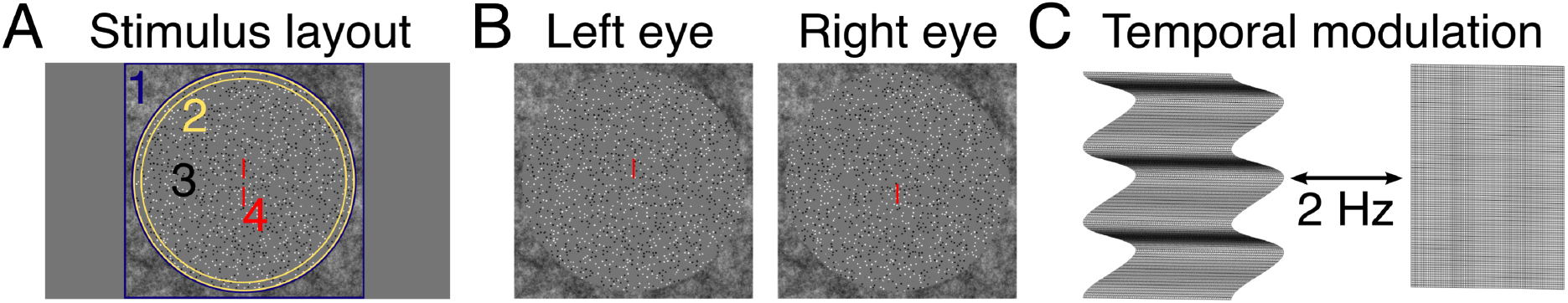
Stimulus details. Panel A shows the general layout common to all stimulus conditions. Stimuli were viewed through a circular aperture embedded within a peripheral 1/f noise fusion lock (1). A ring of decorrelated dots (2) separated the edge of the stimulus. The stimulus was a dynamic random dot stereogram (DRDS, 3) where the corrugation frequency was disparity-defined and followed a sine-wave profile, except in the absolute disparity condition where it was a flat plane. Central nonius lines helped control eye gaze (4). Panel B shows screenshots of the stimulus as viewed by the left and right eyes. Note that in the monocular half-images, the decorrelated dots around the edge are indistinguishable from the DRDS, until they are fused and convey no depth information. Panel B is cross-fusible to reveal a sinusoidal disparity grating. Panel C illustrates the stimulus alternation between a corrugated surface and a flat, zero-disparity plane. The modulation rate was 2 Hz and generated the fundamental frequency in the steady-state visual evoked potential.

Dots within the DRDS were 6 arcmin in diameter and were presented at a density of 15 dots/degree^2^. The placement of the dots was pseudorandom, and to avoid dot overlap we introduced a dot spacing criterion where dots were separated by at least 1.5 x their width. The dot update rate was 20 Hz, meaning that dots were regenerated in new positions on each 6^th^ video frame. Because the frequency of the dot update rate is detectable on the Fourier spectrum of the EEG, we were able to use this as an exclusion criterion against participants who showed weak overall visual evoked responses to the stimulus.

We manipulated the disparity-defined corrugation frequency of the DRDS in 8 separate stimulus conditions for the corrugation tuning experiment, ranging in seven roughly log-spaced steps between 0.1 cpd and 2 cpd with an additional 0 cpd (absolute disparity) condition. At our 70 cm viewing distance each display pixel subtended 10.8 arcsec. Dots in our DRDS stimuli were drawn in OpenGL using anti-aliasing, allowing us to present disparities at sub-pixel resolution. The upper limit for resolving a disparity-defined grating is constrained physically by the resolution limit of the monitor and the size of the dots and biologically by the disparity gradient (Burt and Julesz, 1980; Filippini and Banks, 2009) and underlying receptive field size properties (Banks et al., 2004; Nienborg et al., 2004). The upper limit for the corrugation frequency was near the limit of what could be achieved on our system, given the dot size and spacing constraints that determined the sampling of the disparity surface. Above 2 cpd, the rendering of the dots in the monocular half-image began to look irregular.

The corrugation frequency of the grating was defined in the following manner. First, the DRDS was split into bars of alternating ‘zero-disparity’ and ‘crossed-disparity’ pairs, such that an integer number of bar pairs was viewed though the aperture to achieve a desired corrugation frequency. Second, the overall bar pattern was smoothed to generate sine-wave, rather than square-wave, modulation in depth. This was done by multiplying the magnitude of the dot shift between left and right eye dot pairs that generate the disparity cue with a sine-wave function. The resultant stimulus looks like a 3D wave viewed from above (Figure 1, panel B). The peak of the sine-wave fell in the centre of the crossed-disparity bar, whilst the trough of the sine wave fell in the centre of the zero-disparity bar.

The display alternated in time between a crossed-disparity corrugated surface, and a flat zero-disparity plane (Figure 1, panel C) or in the case of the absolute disparity condition between a flat disparate surface and a zero-disparity surface. The alternation rate was 2 Hz.

The peak-to-trough disparity amplitude of the grating was “swept” in 10 equal log steps over each 10 s stimulus presentation. The stimulus completed two disparate/non-disparate cycles at each step in the disparity sweep. Because disparity sensitivity varies as a function of corrugation frequency, two different sweep ranges were chosen. For ‘low sensitivity’ corrugation frequencies, disparity amplitude was swept between 0.5 and 8 arcmin, whereas for ‘high sensitivity’ conditions the disparity amplitude was swept between 0.2 and 6 arcmin. Low sensitivity conditions were absolute disparity (0 cpd), 0.1 cpd, 1.21 cpd, and 2.00 cpd. High sensitivity conditions were 0.16 cpd, 0.27 cpd, 0.45 cpd, and 0.74 cpd. Optimal sweep ranges were chosen on the basis of pilot experiments, such that the disparity response emerged from the noise in the first half of the sweep and did not saturate towards the end.

### Procedure

For all experiments, trials began with a 1 s prelude in which the display presented the first 60 frames of the upcoming disparity sweep, allowing the EEG and the adaptive filter to reach a steady state. This prelude was followed seamlessly by the 10 s stimulus presentation period, during which disparity amplitude was the swept parameter. The trial ended with a 1 s postlude, recycling the last 60 frames of the stimulus. There was a 2 s gap between subsequent trails, during which participants were instructed to blink as needed. In the reference effects experiment (experiment 1), participants viewed the absolute disparity stimulus with two different tasks superimposed on the disparity modulation in two separate stimulus conditions (see below, ‘Fixation Tasks’). Participants completed 20 trials of each condition, split into presentation blocks of 10 trials each and where each block contained stimuli from one condition. The order of blocks was randomised between participants and breaks were permitted between each block. In total, 20 x 10 s trials were acquired for each of the 2 fixation tasks, per participant. In the corrugation frequency tuning experiment (experiment 2), participants completed 20 trails of each condition, split into blocks of 10 trials each where each block contained stimuli from one of the 8 corrugation frequency conditions. The order of blocks was randomised between participants and breaks were permitted between each block. In total, 20 x 10 s trials were acquired for each of the 8 corrugation frequencies, per participant.

### Fixation Tasks

During stimulus trials, participants were asked to attend to a change at central fixation and press a button to indicate when the change had occurred. The purpose of the task was to encourage fixation at the centre of the screen, allowing convergence on the plane of the display, and to monitor the attentional state of the participants. In the first experiment, we compared responses recorded with two different fixation tasks that varied the availability of a disparity cue from the fixation target. In the first task, a binocularly viewed, intermittently presented letter (either and X or an O) was presented at zero disparity between two nonius lines. The vertically separated nonius lines were presented to the left and right eyes individually and thus did not convey a disparity cue. Participants pressed a button when the X changed to an O. In the second task, the nonius lines themselves changed colour from blue to red and the binocular letters were not presented. Data recorded during the nonius colour task was compared to that obtained in the X-O task in the first experiment and only the nonius task was subsequently used in the spatial tuning experiment. The initial duration of the letter or colour change was 0.5 sec and was varied on a staircase that maintained an 82% correct level of performance.

### EEG Acquisition and Pre-Processing

High-density, 128-channel electroencephalograms (EEG) were recorded using HydroCell electrode arrays and an Electrical Geodesics Net Amps 400 (Electrical Geodesics, Inc., Eugene, OR, USA) amplifier. The EEG was sampled natively at 500 Hz and then resampled at 420 Hz, giving 7 data samples per video frame. The display software provided a digital trigger indicating the start of the trial with millisecond accuracy. The data were filtered using a 0.3 – 50 Hz bandpass filter upon export of the data to custom signal processing software. Artifact rejection was performed in two steps. First, the continuous filtered data were evaluated according to a sample-by-sample thresholding procedure to locate consistently noisy sensors. These channels were replaced by the average of their six nearest spatial neighbours. Once noisy channels were interpolated in this fashion, the EEG was re-referenced from the Cz reference used during the recording to the common average of all sensors. Finally, 1 sec EEG epochs that contained a large percentage of data samples exceeding threshold (30 – 80 microvolts) were excluded on a sensor-by-sensor basis.

### Fourier Decomposition and Filtering

The steady-state VEP (SSVEP) amplitude and phase at the first four harmonics of the disparity update frequency (2 Hz) were calculated by a Recursive Least Squares (RLS) adaptive filter (Tang and Norcia, 1995). The RLS filter consisted of two weights – one for the imaginary and the other for the real coefficient of each frequency of interest. Weights were adjusted to minimise the squared estimation error between the reference and the recorded signal. The memory-length of the filter was 1 s, such that the learned coefficients were averaged over an exponential forgetting function that was equivalent to the duration of one bin of the disparity sweep. Background EEG levels during the recording were derived from the same analysis and were calculated at frequencies 1 Hz above and below the response frequency, e.g., at 1 and 3 Hz for the 2 Hz fundamental. Finally, the Hotelling’s T^2^ statistic (Victor and Mast, 1991) was used to test whether the VEP response was significantly different from zero.

### Dimension Reduction via Reliable Component Analysis

Reliable Components Analysis (RCA) was used to reduce the dimensionality of the sensor data into interpretable, physiologically plausible linear components (Dmochowski et al., 2015). This technique optimizes the weighting of individual electrodes so as to maximize trial-to-trial consistency of the phase-locked SSVEPs. Components were learned on RLS-filtered complex value data, and were learned on the 1F1, 2F1, 3F1 and 4F1 responses across all trials, all participants and all conditions. The Rayleigh quotient of the cross-trial covariance matrix divided by the within-trial covariance matrix was decomposed into a small number of maximally reliable components by solving a generalized eigenvalue problem. Each component can be visualized as a topographic map by weighting the filter weights by a forward model (Haufe et al., 2014; Dmochowski et al., 2015) and yields a complex-valued response spectrum for that component.

Participant-level sensor-space data were weighted by the two most reliable spatial filters, RC1 and RC2. Group-level amplitude and phase estimates for signal (1F1 and 2F1) and noise (side bands of the 1F1 and 2F1 harmonics, respectively) frequencies were calculated by first taking the vector mean across real and imaginary components, across all trials within the same condition. Amplitude was calculated by taking the square root of the sum of the squared real and the squared imaginary components. Phase was calculated by taking the inverse tangent of the real and imaginary components. These vector-averaged amplitude and phase estimates were used to derive neural thresholds (see section below, ‘Estimating neural thresholds’).

For visualizing sweep data, for extracting suprathreshold responses, and for further statistical analyses, we determined the magnitude of the projection of each participant’s response vector on to the group vector average (Hou et al., 2009). Each individual response vector amplitude was multiplied by the cosine of the phase difference between it and the mean vector (Hou et al., 2009). The magnitude of these projections was then used to calculate the group mean projected amplitude and its standard error was calculated using a geometric approach described in detail in (Pei et al., 2017). This procedure yields a group mean amplitude that is very close to the group vector mean, but is advantageous as it converts the vector data to scalar data so that conventional univariate and multi-variate statistics can be used while retaining the improvement in the group-level signal-to-noise ratio when the SSVEP is phase consistent across subjects. The measure also minimizes error estimates that are non-normally distributed.

### Estimating Neural Thresholds

Because 1F responses increase linearly with log disparity amplitude (Norcia et al., 1985b) a linear function was fit to the group-level disparity response functions to estimate a neural threshold for each corrugation frequency as the zero-amplitude intercept (Campbell and Maffei, 1970; Norcia et al., 1985a). Our fitting function searched for a range of at least two consecutive 1 s bins where amplitude was both monotonically increasing and likely dominated by signal and thus usable for extrapolation to zero amplitude. The range was established on the basis of the following criteria. First, to avoid fitting spikes in the record caused by artifacts, the amplitude at the noise frequencies in a bin could not exceed 70% of the amplitude of the signal in a bin. Second, the *p* value of a the Hotelling’s *T*^2^ test was below 0.160 (at least 1.5 standard deviations from zero). Third, the noise in the frequency side bands did not exceed 30% of the signal in the same bin. Fourth, the phase difference compared to the previous bin was not greater than −100 and smaller than +80 minimizing fitting over non-physiological data where the response phase lags with increasing visibility. Fifth, the amplitude was monotonically increasing, and the signal was larger than that measured in the previous bin. Finally, the SNR of the signal was greater than 1.5. Bins that satisfied these criteria were deemed likely to contain meaningful signals, and a linear function was fit to consecutive bins that satisfied these conditions. The x-axis intercept was taken as the neural threshold – the disparity at which the cortical response would have been zero in the absence of additive EEG noise. In some cases, especially when amplitudes were low and changes in amplitude from bin-to-bin were small, the slope of the fit was shallow resulting in an over-extrapolated threshold. This was deemed to be the case when there were more than two bins between the first bin used to fit the function, and the estimated threshold. When this occurred, the threshold was set to the disparity value at the bin prior to the first bin containing measurable signal. This occurred in one instance, for the neural threshold of the 1F1 response in RC1, in the 0.10 cpd condition. An example of the scoring procedure is shown in Figure 5, panel B.

We constructed a Disparity Sensitivity Function (DSF) from the estimated neural thresholds. To compare the shape of the ‘neural DSF’ against the DSF measured in a range of psychophysical studies (Figure 5, Panel D: (Tyler, 1973; Schumer and Ganz, 1979; Pulliam, 1982; Rogers and Graham, 1982; Lankheet and Lennie, 1996; Lee and Rogers, 1997; Bradshaw and Rogers, 1999; Hess et al., 1999; Hogervorst et al., 2000; Tyler and Kontsevich, 2001; Bradshaw et al., 2006; Serrano-Pedraza and Read, 2010; Didyk et al., 2011; Kane et al., 2014; Peterzell et al., 2017)), we extracted reported data using WebPlotDigitizer (software freely available at https://automeris.io/WebPlotDigitizer/) and normalized by dividing each threshold against the lowest threshold in each dataset, forcing each DSF to bottom out at 1. Where both upper and lower limits of disparity sensitivity were measured, the upper limit datapoints were excluded.

### Suprathreshold Disparity Tuning Functions

Disparity tuning functions were also estimated from the projected amplitude data from suprathreshold disparities to maximize the signal to noise ratio and to allow comparisons to be made across different conditions using conventional scalar-valued statistics. Because different sweep ranges and different disparity-step values were used in different corrugation frequency conditions, we used linear interpolation to estimate signal amplitude at 6 and 2 arcmin. Estimates were generated for each participant using their mean projected amplitudes calculated across all trials, giving a sweep function from which the amplitudes at the two disparity levels were estimated.

### Statistical Analyses

Suprathreshold data were analysed using repeated measures analysis of variance (ANOVA) in R, using functions from the *rstatix* package (in particular, anova_test which is a wrapper for car::Anova). Data were assessed for normality using the Shapiro-Wilk test and visual inspection of QQ plots. Sphericity was assessed using Mauchly’s test. Where sphericity was violated, the Greenhouse-Geisser correction was applied. The generalized effect size (Olejnik and Algina, 2003), which estimates the proportion of variability explained by the within-subjects factor, is reported with each *F* test. Qualitative descriptors of effect size are consistent with Cohen’s benchmarks, with small, medium and large effects ascribed to effect sizes of 0.2, 0.5 and 0.8, respectively (Olejnik and Algina, 2003). Pairwise *t*-tests were used to interrogate main effects and interactions and reported *p*-values are adjusted for multiple comparisons using the Bonferroni correction.

## RESULTS

### Topography of the Disparity Response

Reliable component analysis (RCA) was used to identify a small number of disparity-specific sources via differences in their spatial topography and trial-to-trial consistency. Performing RCA on our data revealed two primary response components that had statistically reliable responses to changing disparity across all corrugation frequencies and at both response frequencies of interest, 1F1 and 2F1. The topography of RC1 was focused over midline occipital electrodes, whereas RC2 was right-lateralized over occipito-temporal areas and was more spatially diffuse (Figure 2). These components are similar to those identified in an earlier study of changing disparity responses generated by DRDS (Norcia et al., 2017). Statistically reliable first and second harmonics of the disparity update frequency 1F1 and 1F2 were present consistently in both RC1 and RC2. Subsequent analyses therefore focused on 1F1 and 2F1 signals in RC1 and RC2.

**Figure 2:**
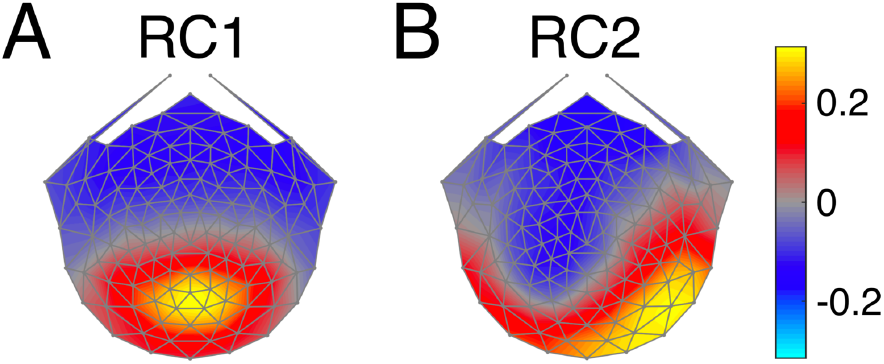
Topographies of the two most consistent neural responses associated with the stimulus disparity update, extracted via Reliable Component Analysis learned on the corrugation frequency tuning data. In RC1 (Panel A), electrodes over midline occipital areas were weighted more strongly. In RC2 (Panel B), the response was right-lateralized over occipito-temporal areas. Heat maps represent the weighting assigned to each electrode, but the sign of the weight is arbitrary.

### Eliminating Reference Effects in the Absolute Disparity Stimulus

Fixation targets can create unwanted reference effects when trying to estimate responses to absolute disparity and the absence of binocular references is critical for generating a nearly ‘pure’ absolute disparity stimulus. In our first experiment, we therefore measured the possible effect of fixation point references used to control vergence. We compared a binocular central fixation task that created a relative disparity reference to a dichoptic nonius line fixation task did not. In the former case, a binocularly viewed changing letter was presented centrally at zero disparity half-way between two nonius lines. For the monocular fixation task, the nonius lines themselves changed colour and there was no binocular reference. In this condition, the only references available come from the peripheral fusion-lock and monitor bezel.

The effect of the presence of a binocular reference is in shown in Figure 3. In both RC1 (Panel A) and RC2 (Panel B), there was a robust 1F1 response to the disparity change when there was a binocular reference at fixation. However, there was a marked *reduction* in the amplitude of the 1F1 response when the task was switched from the binocular X-O task to the nonius colour-change task. Within each RC, the amplitude of the responses between conditions were compared using paired-samples *t*-tests at each bin, and bins where this difference was significant (*p* < .050) are marked in Figure 3.

**Figure 3:**
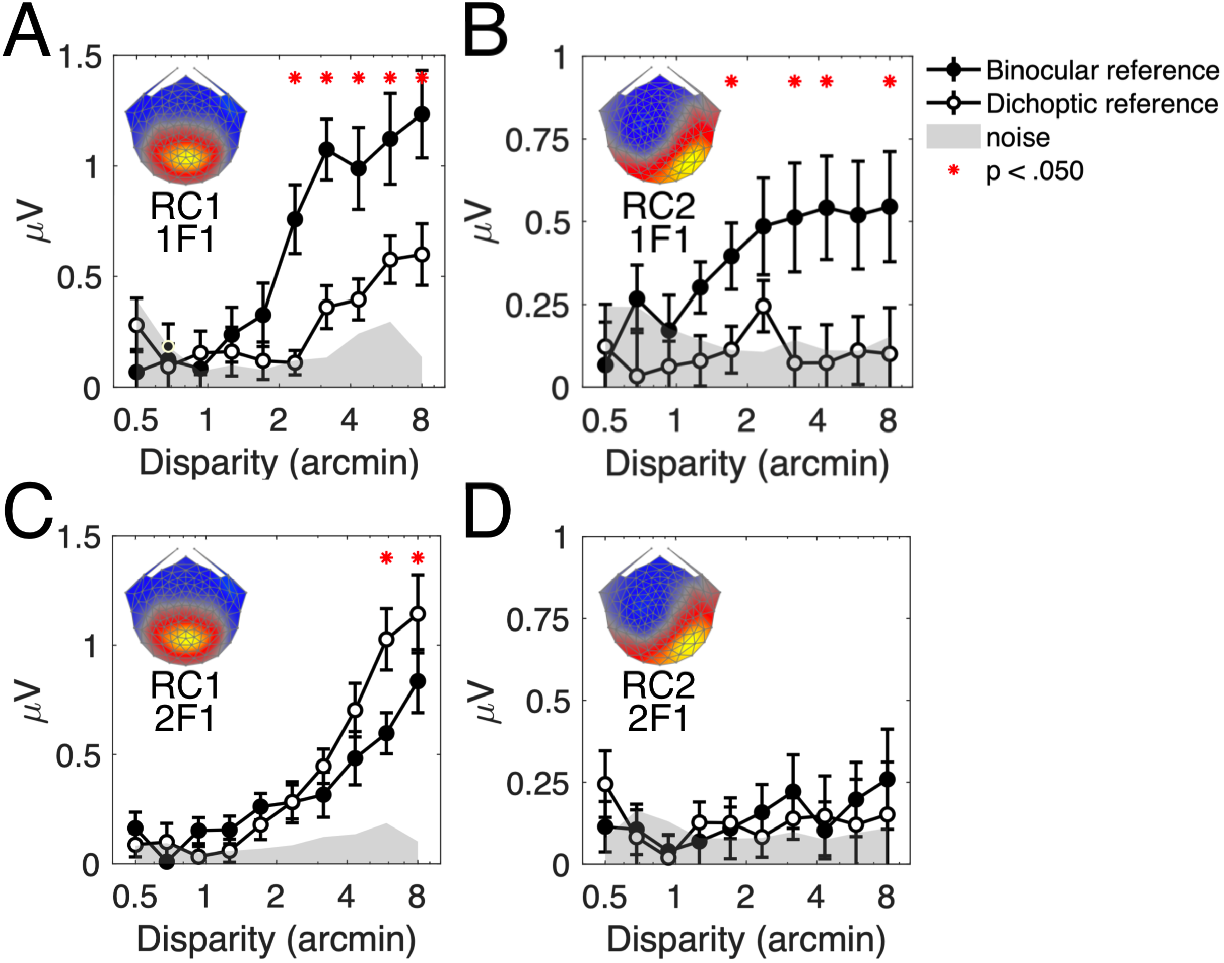
Effect of a central fixation task on the 1F1 and 2F1 response amplitudes. Responses were evoked by an absolute disparity stimulus oscillating between zero disparity and a given crossed disparity. In all panels, the amplitude of the evoked response is indicated as a function of disparity. The top row (panels A and B) shows the 1F1 response weighted by RC1 and RC2, respectively. The bottom row (panels C and D) shows the 2F1 response weighted by the same RCs. RC weightings are illustrated in the topography map inserts in each panel. Participants performed an attention task placed on the nonius lines (‘dichoptic reference’, open circles) or a letter change task (‘binocular reference’, filled circles). The introduction of a binocular reference adds a zero-disparity reference to the foveated part of the stimulus, introducing a relative disparity signal into the otherwise absolute disparity stimulus. Thus, the 1F1 response component is sensitive to relative disparity, especially in RC1, whilst the 2F1 response is unperturbed. RC2 is insensitive to pure absolute disparity, and the lack of a 2F1 response in this RC implies that 2F1 is driven primarily by absolute disparity.

For RC1, the 1F1 sweep response during the dichoptic reference condition was shifted rightwards along the x-axis, indicating that the response emerged only at higher disparity amplitudes. This residual 1F1 response may be due to some relative disparity leakage arising from imperfect separation of the edge of the changing disparity region and the static zero-disparity fusion lock or monitor bezel. Alternatively, a functional explanation could be that the 1F1 signal is driven by asymmetries in the preferred direction of motion in depth (Cottereau et al., 2011). In contrast, in RC2, the amplitude of the 1F1 response during this condition rarely rose above the noise floor.

The sensitivity of the VEP to a binocular reference at fixation demonstrates that the asymmetric 1F1 response is highly sensitive to the presence of more than one disparity in the stimulus. This result is consistent with previous results (Cottereau et al., 2012b) where the dynamic nature of the changing disparity stimulus also resulted in ‘making and breaking’ of the disparity plane defined by a binocular zero disparity reference. Thus, the 1F1 response is a strong readout for relative disparity mechanisms. The small residual magnitude of this response in the nonius fixation condition indicates that the response is dominated by contributions from absolute vs. relative disparity mechanisms.

The 2F1 response in RC1 (panels C and D) is strongly dependent on disparity, but is comparatively less perturbed by the change in disparity reference. In contrast to the 1F1 response, the 2F1 amplitude is somewhat higher during the ‘pure’ absolute disparity condition especially at large disparities. This implies that it is strongly driven by absolute disparity modulation. In RC2 (panel D), the second harmonic fails to evoke a strong response for either the binocular or the monocular conditions, implying that the sources underlying RC2 are relatively insensitive to absolute disparity modulation.

### Associating 1F1 and 2F1 Responses with Sustained and Transient Activity

To distinguish temporal response components that are reflective of transient vs. sustained mechanisms, we used a spectral analysis approach previously introduced for the study of contrast evoked potentials (McKeefry et al., 1996). McKeefry et al. used simulations to argue that sustained responses like those we observe for the stereo grating will manifest in the first harmonic of the SSVEP, whereas transient mechanisms manifest in the even harmonics.

This approach applied to our disparity-driven responses is illustrated in Figure 4, which shows spectra (leftmost panels) and time courses (middle and rightmost panels) of the responses to a flat plane with nonius lines at fixation (dichoptic reference, top row) vs. the XO task at fixation (binocular reference, bottom row). Data are taken from the experiment described above, and are single-cycle group averages across a cluster of midline occipital electrodes that underlie RC1 (71, 72, and 76).

**Figure 4:**
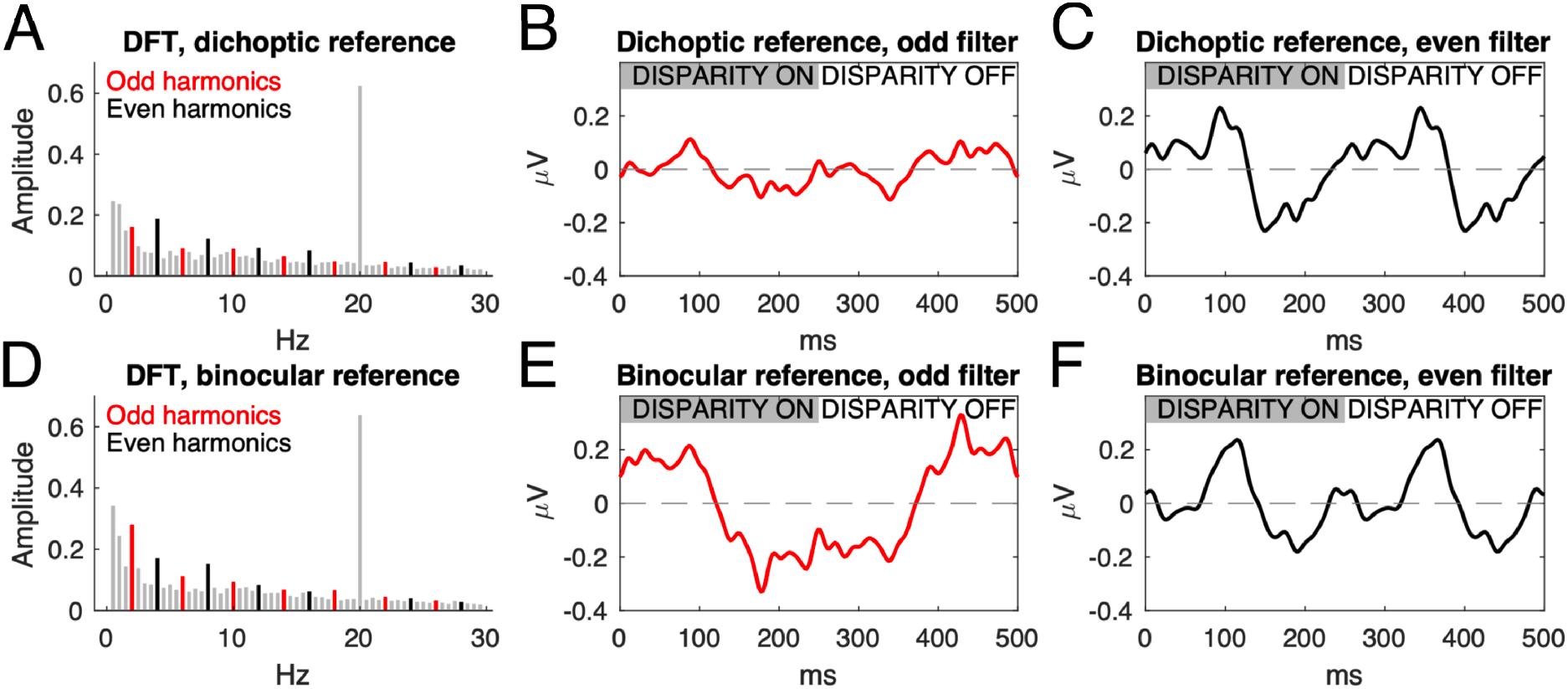
Harmonic-based filtering approaches reveal the sustained and transient response components of the SSVEP. Participants viewed a disparity plane alternating between crossed disparity (first 150 ms, indicated by grey bar in middle and right columns) and zero disparity. Fixation was encouraged using nonius lines (dichoptic reference, no relative disparity signals) or an X-O task at fixation (binocular reference, inducing relative disparity cues). Filtering on the odd (in red) or even (in black) harmonics reveals sustained and transient response components in the reconstructed waveforms, respectively (middle and right columns). Gray bars in the discrete Fourier transforms (left column) are frequencies that are nulled by the filtering – the large 20 Hz dot update response has also been removed. The introduction of a binocular reference point at fixation results in a sustained negative-going potential, shifted ~120 ms relative to stimulus onset, that is revealed by the odd filter, which is absent in the pure absolute disparity condition when only nonius lines are present. Both conditions have transient response components revealed by the even filter. Data are group-level averages of single-cycle responses across all trials, for a cluster of midline occipital electrodes.

The simple addition of a binocular reference at the fovea dramatically alters the EEG waveform, particularly apparent under the ‘odd’ filtering regime (middle column). Reconstructing the response to the binocular reference condition using only the odd harmonics (1F1, 3F1, 5F1 and so on) of the fundamental frequency (2 Hz) shows a nearly square-wave sustained response. Reconstructing the response from the dichoptic reference disparity plane condition using only the even harmonics (2F1, 4F1, 6F1 and so on) yields a biphasic response after both onset and offset of the stimulus. The filtering approach is illustrated in the leftmost panels, where odd harmonics are in red and ever harmonics are in black. Nulled frequencies are plotted in grey. By splitting the response into its odd and even harmonic components, we emphasize the sustained activity in the (low-order) odd harmonics and transient activity in the even harmonics. In practice, odd and even harmonic responses above the 1^st^ and 2^nd^ harmonics are small and difficult to measure over a wide range of stimulus conditions, so in the remainder we focus on 1F1 and 2F1 as correlates of sustained and transient activity.

### Properties of the 1F1 Response in RC1

#### Disparity response functions vary with corrugation frequency

If the 1F1 response is an indicator of relative disparity processing, it should be tuned for corrugation frequency. In particular, the response to disparity gratings that contain multiple disparities and therefore relative disparity should differ from the flat plane, absolute disparity condition that does not. We thus measured evoked responses while sweeping the amplitude of disparity-defined gratings that varied in their corrugation frequency. Because the stimulus modulated between zero disparity and crossed disparity, evoked responses locked to changing disparity could be generated in two ways: first, from the local change in disparity within a single temporal stimulus cycle (e.g., from changes in local, absolute disparity) or secondly, from the change from a flat plane at fixation, to a corrugated surface in depth (relative disparity). The *magnitude* of the 1F1 response was found to increase with the disparity amplitude. We first analysed this sweep response in the most reliable component (RC1). Sweep responses for all stimulus conditions are overlaid on the same axis in Figure 5, panel A. The lateral shift in the responses along the x-axis implies that sensitivity to disparity changes as a function of corrugation frequency. Note that the conditions with the earliest, and largest, responses above the noise floor lie in the mid-ranges of the corrugation frequencies we tested. Least sensitive are the responses to very high or very low corrugation frequencies, with the weakest response to the absolute disparity stimulus. In the absolute disparity condition, the signal emerged from the noise relatively late in the sweep and the response was more than a factor of three weaker as compared to more sensitive conditions.

**Figure 5:**
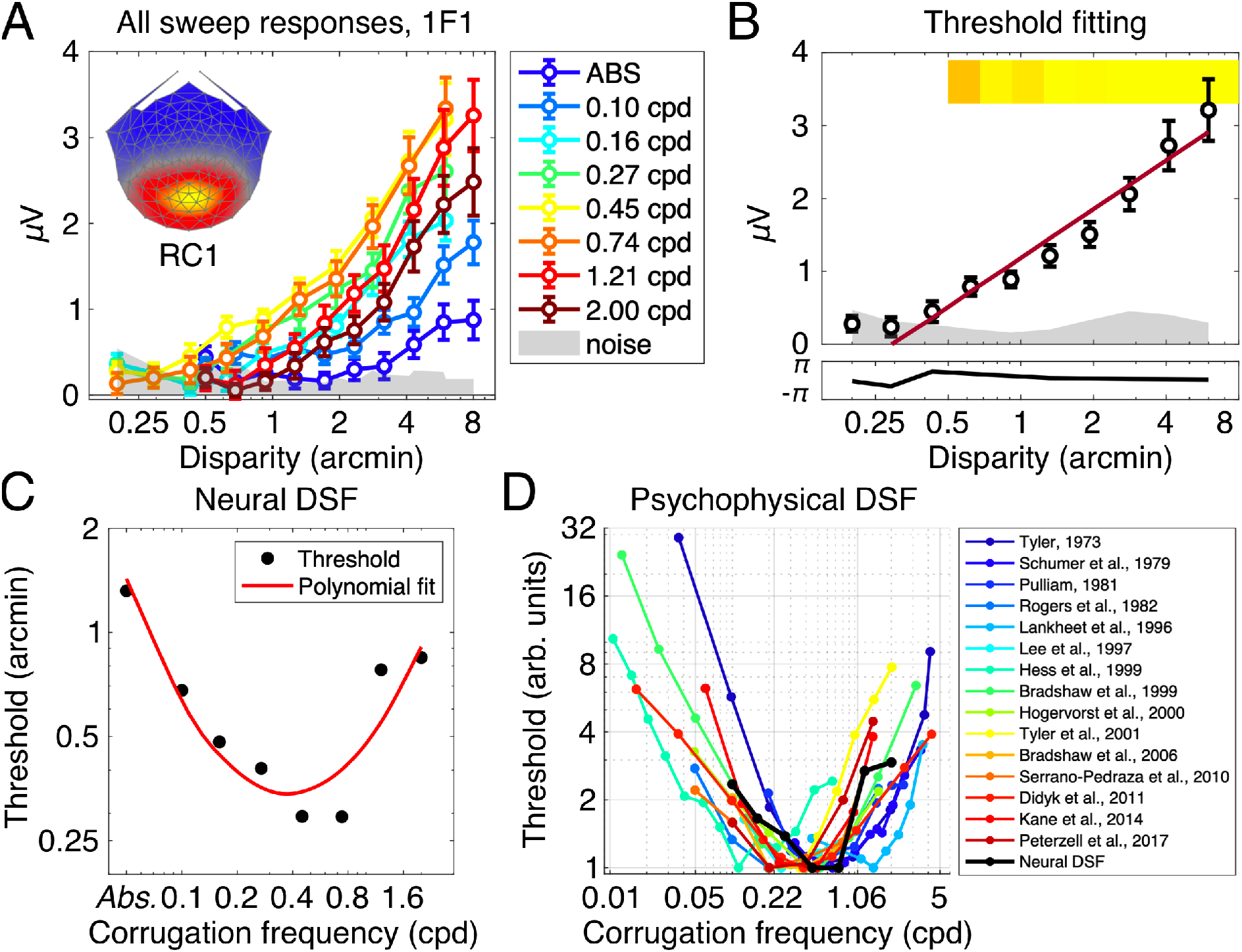
Results from the 1F1 response in RC1 showing dependence on corrugation frequency. Panel A shows all sweep responses, plotted as the amplitude of the 1F1 response against the peak-to-trough disparity amplitude of the grating. Different colours denote different corrugation frequency conditions, and all error bars are ±1SEM. The shaded area is the mean noise amplitude in the frequency side bands, averaged across conditions. Weightings for RC1 at each electrode are shown in the topographical map insert. Panel B shows an example of the neural threshold estimation, here, data from the 0.45 cpd condition are shown. A linear trend (in red) is fit against bins that fulfil a set of criteria (see Methods for more details). The intersection of the linear trend with the x-axis is taken as the estimated neural threshold. Thresholds for each condition are plotted in Panel C and demonstrate U-shaped corrugation frequency sensitivity. The corrugation tuning in C is consistent with previous psychophysical measurements of the disparity sensitivity function, shown in Panel D (y-axis is normalized, with the best threshold measured for each study being set to 1. ‘Absolute disparity’ condition in Neural DSF data omitted for consistency on the x-axis).

#### The disparity threshold for 1F1/RC1 is tuned for corrugation frequency

Because the magnitude of the disparity amplitude increases monotonically and is approximately linear, it is possible to estimate a neural threshold by regression to zero amplitude of the disparity response function (Norcia et al., 1985a). The value at which the linear function crosses the x-axis is taken as the neural threshold and is indicative of the smallest disparity required to elicit a systematic neural response. An example of this process is illustrated in Figure 5, panel B.

We extracted neural thresholds for 8 different corrugation frequency conditions and found that they form a U-shaped function of corrugation frequency tuning (Figure 5, panel C). In both our measurements and previous psychophysical ones (Figure 5, panel D), disparity sensitivity is maximal between ~ 0.5 and 0.75 cpd. In the limiting case (absolute disparity, or 0 cpd), the threshold we measure is on the order of four times higher than at peak sensitivity. Our monitor resolution and rendering capabilities limited the maximum usable corrugation frequency to 2 cpd where the threshold (0.85 arcmin) was around three times higher than that measured at peak sensitivity.

#### Suprathreshold response amplitude mirrors the Disparity Sensitivity Function

The disparity response functions of Figure 6, panel A each increase monotonically with disparity amplitude and have a similar slope. This implies that the shape of the corrugation tuning should be independent of disparity amplitude over the supra-threshold disparity range. To test this quantitatively, we plotted log evoked response amplitude as a function of corrugation frequency for two supra-threshold disparities (Figure 5, panel A). Because the sweep ranges varied across condition, we extracted interpolated amplitudes for 6 and 2 arcmin measurements. At both disparity amplitudes, the tuning function is similar to an inverted DSF (Figure 5, Panel C), where the response amplitude peaks between 0.45 and 0.74 cpd.

**Figure 6:**
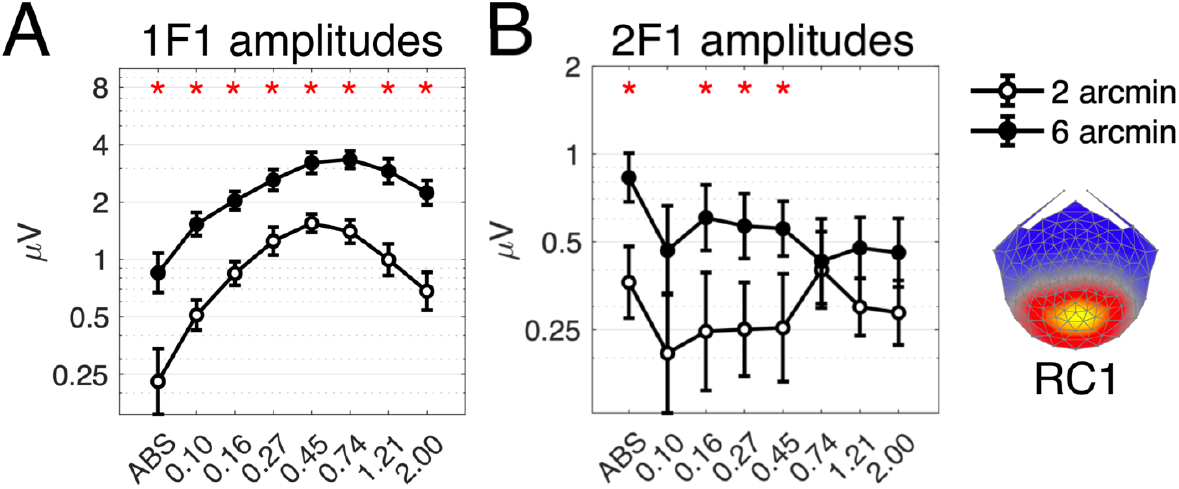
Suprathreshold response amplitudes for different corrugation frequency conditions in RC1, at two different disparity amplitudes (2 arcmin, open circles; 6 arcmin, filled circles). The 1F1 response is shown in Panel A. The 2F1 response is shown in Panel B. The asterisk above each bin shows amplitudes that are significantly different (with Bonferroni correction) between the two disparity levels, as calculated by pairwise comparisons after the ANOVA. Error bars are ±1SEM and the topographical map insert illustrates the weightings across electrodes applied for calculating the mean response in RC1.

To formally quantify the effects of disparity level, corrugation frequency, and interactions between these, we performed a two-way repeated measures ANOVA on the data plotted in Figure 6, panel A. The majority of the variance in the data was captured by the main effect of disparity level (*F*(1, 24) = 63.59, *p* < .001, generalized effect size (*η^2^_G_*) = 0.23) where the mean amplitude of the responses at 6 arcmin was significantly larger than the mean amplitude at 2 arcmin. The main effect of corrugation frequency was also highly significant *F*(2.57, 61.61) = 17.93, *p* < .001, *η^2^_G_* = 0.18, with Greenhouse-Geisser correction for sphericity as assessed by Mauchly’s test (*W* = 0.01, *p* < .001)), indicating that, as expected, the amplitude of the 1F1 response component was dependent on the corrugation frequency of the DRDS stimulus.

We also measured a significant interaction between disparity level and corrugation frequency *F*(3.93, 94.29) = 6.16, *p* < .001, *η^2^_G_* = 0.03, with Greenhouse-Geisser correction for sphericity as assessed by Mauchly’s test (*W* = 0.07, *p* < .001)). This interaction was driven by the slight shift in the peak of the function as well as the more exaggerated U-shape at 2 arcmin, where the amplitude of the response increased almost seven-fold (as opposed to four-fold at 6 arcmin) from absolute disparity to the peak of the function. However, we think it unlikely that the neural sources of the 1f1 response are different at 6 arcmin than at 2 arcmin, as is indicated by the high degree of correlation (*R*(14) = .94, *p* < .001) between the two functions as well as the small effect size of the interaction.

### Properties of the 2F1 Response in RC1

#### Sweep functions for 2F1/RC1 reveal little tuning

All sweep responses captured by the 2F1 response in RC1 are plotted in Figure 7, Panel A. Although we measured reliable sweep responses in all conditions, we observed that many of the responses overlapped across different corrugation frequency conditions. Notably, the response to the *absolute* disparity stimulus was now the largest, where for the 1F1 response it was the weakest. Thus, though we observe little evidence for corrugation frequency tuning in the 2F1 responses, we note its response to absolute disparity is particularly robust.

**Figure 7:**
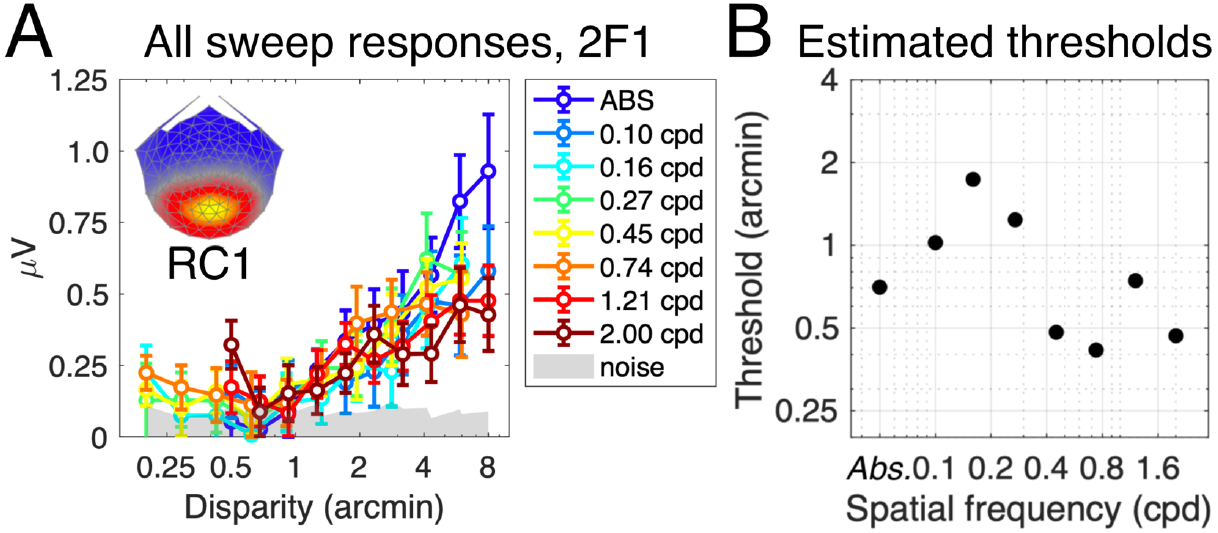
Results from the 2F1 response to disparity in RC1. Responses to gratings at different corrugation frequencies are plotted in Panel A, where the signal increased in amplitude as the peak-to-trough disparity in the grating increased. Neural thresholds were extracted by fitting a linear function to each curve in A. The value of the x-axis intercept was taken as the threshold, and is plotted in Panel B. Both panels indicate no systematic tuning to corrugation frequency, in contrast to the 1F1 response which was strongly tuned. Note the sensitivity of the 2F1 response to absolute disparity, expressed as a large signal amplitude in Panel A (dark blue curve) and a relatively low threshold in Panel B (leftmost point on the plot).

#### Lack of corrugation frequency tuning at 2F1/RC1 threshold

We also estimated 2F1 neural thresholds in RC1 for each corrugation frequency condition using the same method as for the 1F1 responses. Results are plotted in Figure 7, Panel B. The 2F1 response did not show the same systematic tuning to corrugation frequency as the 1F1 response, where thresholds followed the U-shaped function of the DSF. Instead, the estimated 2F1 thresholds are irregularly scattered between 0.4 and 1.7 arcmin, indicating little tuning to corrugation frequency.

Instead, we observed that the response to *absolute* disparity was particularly sensitive, with a large response amplitude in comparison to the other corrugation frequency conditions (blue curve in Figure 7, panel A). In addition, the extracted threshold for the 2F1 absolute disparity condition was markedly lower than for the simultaneously recorded 1F1 response in the same condition (0.70 vs. 1.32, respectively). Whilst the 2F1 response in RC1 is not particularly sensitive to corrugation frequency – and therefore to relative disparity – it does appear to be a stronger readout for the absolute disparity mechanism.

#### Suprathreshold responses are untuned in 2F1/RC1

Similar to the 1F1 analysis, we investigated the effects of disparity amplitude and corrugation frequency on the amplitude of the 2F1 responses potted in Figure 6, panel B, using a two-way repeated measures ANOVA. Again, there was a main effect of disparity level though the effect size was modest (*F*(1, 24) = 11.04, *p* = .003, *η^2^_G_* = 0.04). Pairwise comparisons revealed that the amplitude of the response at 6 arcmin was greater than at 4 arcmin at absolute disparity, 0.16 cpd, 0.27 cpd and 0.45 cpd, but nowhere else (adjusted *p* = .003, .004, .007 and .017 respectively, Bonferroni corrected for multiple comparisons).

Importantly, there was *no* effect of corrugation frequency, confirming the lack of corrugation tuning in the 2F1 response (*F*(7, 168) = 1.40, *p* = .210, *η^2^_G_* = 0.01). This is in contrast to the 1F1 result, where the main effect of corrugation frequency was highly significant, reflecting a deviation in the corrugation sensitivity of these two harmonics and the neural mechanisms that underpin them. The interaction between disparity level and corrugation frequency was also nonsignificant, indicating that the lack of tuning was consistent across both disparity levels (*F*(3.93, 94.29) = 6.16, *p* < .001, *η^2^_G_* = 0.01).

#### Differential tuning for suprathreshold 1F1 and 2F1 responses in RC1

To directly compare the corrugation tuning properties of 1F1 and 2F1 responses in RC1, we took advantage of the greater signal to noise ratio in the suprathreshold responses extracted at 6 arcmin disparity. 1F1 and 2F1 response amplitudes as a function of corrugation frequency are plotted in Figure 6. The 1F1 response is U-shaped as a function of corrugation frequency for both disparities, whilst the 2F1 response is relatively flat across all corrugation frequencies with the exception of the response to absolute disparity.

We used a two-way repeated measures ANOVA to quantify the effects of harmonic and corrugation frequency on response amplitudes at 6 arcmin, and measured a significant main effect of harmonic (*F*(1, 24) = 78.65, *p* < .001, generalized effect size (*η^2^_G_*) = 0.34) and a smaller significant effect of corrugation frequency (*F*(2.61, 62.57) = 11.80, *p* < .001, *η^2^_G_*= 0.08, with Greenhouse-Geisser correction for sphericity as assessed by Mauchly’s test (*W* = 0.01, *p* < .001)).

The most telling result was the interaction between harmonic and corrugation frequency, which was significant (*F*(2.74, 65.70) = 16.65, *p* < .001, *η^2^_G_* = 0.11, with Greenhouse-Geisser correction for sphericity as assessed by Mauchly’s test (*W* = 0.02, *p* < .001)). This implies that the tuning functions are different between 1F1 and 2F1, where 1F1 shows a dependence on corrugation frequency whereas the 2F1 response does not.

Furthermore, pairwise comparisons showed that the 1F1 response amplitude was greater than the 2f1 response amplitude at all corrugation frequencies except absolute disparity, after Bonferroni correction for multiple comparisons (adjusted *p* values were .932, and < .001 for absolute disparity and all other corrugation frequencies). This echoes the observation that the 2F1 response to absolute disparity is particularly large.

### Properties of the 1F1 response in RC2

#### 1F1 sweep responses show evidence of corrugation frequency tuning

Reliable component analysis allowed us to distinguish a second reliable component, RC2 that was right-lateralized over occipito-temporal electrodes (see Figure 2). This source was sensitive to the presence of a binocular reference at fixation (see Figure 3, panel B) and is thus also sensitive to relative disparity. The same analyses as in RC1 were therefore carried out on the 1F1 signal in RC2. All sweep responses are shown in Figure 8. Sweep responses followed the same pattern as in RC1, where the 0.74 cpd grating yielded the highest response amplitudes and the weakest response was to the 0.1 cpd grating. In contrast to RC1, and the absolute disparity stimulus failed to evoke a response above the noise floor. This may be due to the overall lower SNR in RC2 – in RC1, the largest response to absolute disparity occurred at the top of the sweep (8 arcmin) and its amplitude was a factor of ~4.5 times greater than the noise floor at that disparity amplitude. With at least a 3-fold overall reduction in SNR in RC2, the absolute disparity signal may have been too weak to be reliably measured. It is possible that increasing the maximum disparity of the sweep could evoke a weak 1F1 response to absolute disparity in RC2.

**Figure 8:**
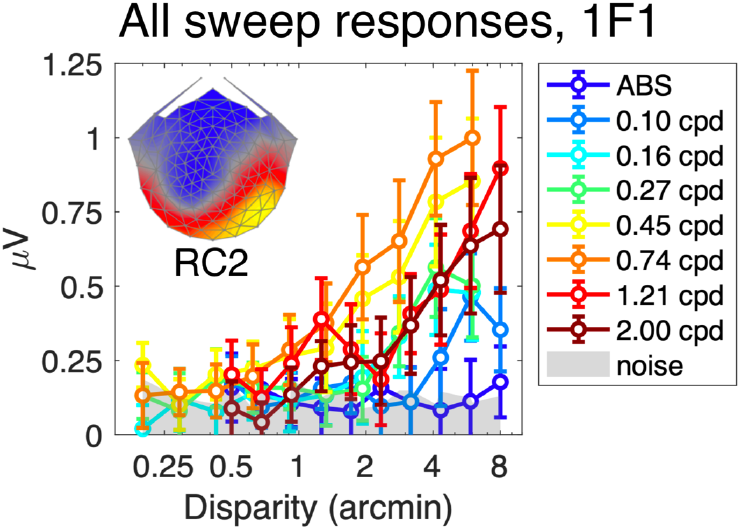
All sweep functions of the 1F1 response in RC2. Response amplitude increases systematically with increasing disparity and vary systematically as a function of the corrugation frequency of the stimulus. Error bars are ±1 SEM and the noise floor is indicated in light grey.

RC2 shows a similar lateral shift in the sweep functions as RC1, indicating tuning to corrugation frequency. 1F1 responses varied systematically as a function of the stimulus corrugation frequency. Compared to RC1, this lateral shift was not as clearly organized, perhaps commensurate with the lower overall SNR.

#### Suprathreshold 1F1 responses are tuned for corrugation frequency

Due to the lower SNR of the RC2 response, we did not estimate thresholds, but rather characterized corrugation tuning from the suprathreshold response functions measured at 2 and 6 arcmin of disparity. Results are shown in Figure 9, Panel A. Here, it is possible to see a peak in sensitivity around 0.74 cpd, and that sensitivity drops off for low and high corrugation frequencies. The signal at 6 arcmin is larger than at 2 arcmin, where the signal is only significantly greater than zero at the peak of the tuning function. Note that the amplitude of the 0 cpd, absolute disparity condition is not different from zero at either disparity amplitude, reflecting the sweep values showing that the signal is weak and barely rises above the noise floor (dark blue line in Figure 8).

**Figure 9:**
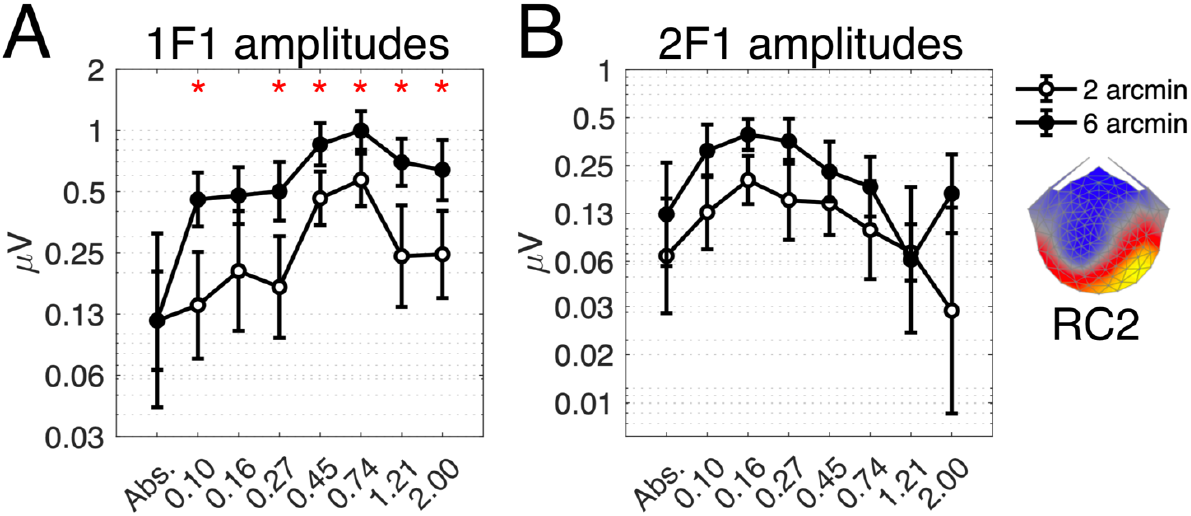
Suprathreshold response amplitudes for different corrugation frequency conditions in RC2, at two different disparity amplitudes (2 arcmin, open circles; 6 arcmin, filled circles). The 1F1 response is shown in Panel A. The 2F1 response is shown in Panel B. The asterisk above each bin shows amplitudes that are significantly different (with Bonferroni correction) between the two disparity levels, as calculated by pairwise comparisons after the ANOVA. Error bars are ±1SEM and the topographical map insert illustrates the weightings across electrodes applied for calculating the mean response in RC2.

We duplicated the analysis of RC1, using a two-way repeated measures ANOVA to measure effects of disparity level and corrugation frequency on response amplitudes of the 1F1 response in RC2. Results echo those in RC1, although the effects were smaller. There was again a significant main effect of disparity level (*F*(1, 24) = 26.67, *p* < .001, generalized effect size (*η^2^_G_*) = 0.04), where amplitude of 6 arcmin response was significantly larger than the 2 arcmin response at all corrugation frequencies except absolute disparity and 0.16 cpd (adjusted *p* values .999 and .088 respectively, Greenhouse-Geisser correction applied).

Importantly, the response varied significantly as a function of corrugation frequency, illustrating that the 1F1 response maintains its corrugation tuning properties in RC2. The main effect of corrugation frequency was significant (*F*(2.31, 55.35) = 4.27, *p* < .001, *η^2^_G_* = 0.06, with Greenhouse-Geisser correction for sphericity as assessed by Mauchly’s test (*W* = 0.00, *p* < .001)), and the effect sizes imply that majority of the variance is explained by the corrugation frequency (*η^2^_G_* = 0.06), not the disparity level (*η^2^_G_* = 0.04).

In RC2, the Interaction between disparity level and corrugation frequency was nonsignificant (*F*(7, 168) = 1.14, *p* = 344, *η^2^_G_* = 0.01, with sphericity assumed as assessed by Mauchly’s test (*W* = 0.16, *p* = .058)) indicating that the 2 arcmin and 6 arcmin measurements followed same tuning profile.

### Properties of the 2F1 Response in RC2

#### Low sensitivity to disparity

Sweep responses in RC2 are shown in Figure 10. In RC2, the sweep responses were noisy and weak, and were generally overlapping, with the clearest responses to stimuli between 0.10 and 0.27 cpd. The two highest corrugation frequency conditions only generated responses above the noise floor in the last two disparity bins. In contrast to RC1, the 2F1 response to absolute disparity did not consistently emerge from the noise floor – and was near zero at the highest disparity level tested. Recall that in RC1, the 2F1 response showed little corrugation frequency tuning and sweep responses were largely overlapping, with the exception of the response to absolute disparity (Figure 9, Panel A). Overall, in RC2 there was at least a factor of 2 reduction in the 2F1 response as compared to the 1F1 response for this component.

**Figure 10:**
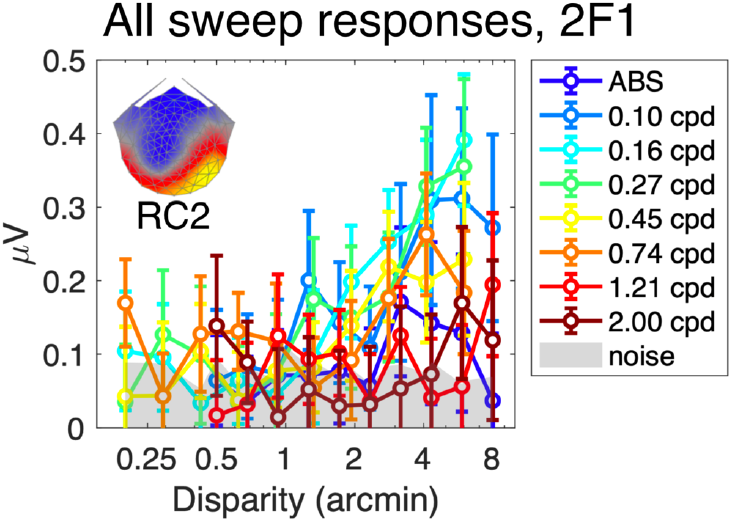
Sweep functions of the 2F1 response in RC2. Response amplitude increases with increasing disparity but the variance over corrugation frequency is less clear. Error bars are ±1 SEM and the noise floor is indicated in light grey.

#### Suprathreshold 2F1 response is untuned for corrugation frequency

Responses at higher disparity amplitudes were extracted to interrogate the tuning function of the 2F1 response in RC2 (Figure 9, Panel B). Here, the peak of the function was shifted towards the lower corrugation frequencies, where the highest response amplitude was measured at 0.27 cpd. The signal was consistently weak across different corrugation frequency conditions, and a higher disparity amplitude (6 arcmin, rather than 2) was required to measure responses that were consistently significantly different from zero. The two highest corrugation frequency conditions, as well as the absolute disparity condition, did not evoke responses that were significant.

Because the signal was low, the two-way repeated measures ANOVA measured only a small significant effect of disparity level (*F*(1, 24) = 7.95, *p* = .010, *η^2^_G_* = 0.02). There was no main effect of corrugation frequency (*F*(4.58, 109.85) = 2.02, *p* < .087, *η^2^_G_* = 0.03 with Greenhouse-Geisser correction for sphericity as assessed by Mauchly’s test (*W* = 0.10, *p* =.006)), and no pairwise comparisons passed the corrected Bonferroni threshold for significance. The interaction was also nonsignificant. Therefore, we do not find compelling evidence for corrugation frequency tuning in the 2F1 component in RC2.

#### Comparing suprathreshold tuning functions for 1F1 and 2F1 in RC2

To compare results across harmonics, compared the 1F1 and 2F1 results at 6 arcmin in RC2 using a repeated measures ANOVA (Figure 9, data in panels A and B). The main effects of harmonic (*F*(1, 24) = 9.38, *p* = .005, *η^2^_G_* = 0.06) and corrugation frequency (*F*(3.42, 81.98) = 2.79, *p* = .044, *η^2^_G_* = 0.03 with Greenhouse-Geisser correction for sphericity as assessed by Mauchly’s test (*W* = 0.05, *p* < .001)) were significant. Pairwise comparisons showed that 1F1 response was larger than the 2F1 response at higher corrugation frequencies (0.45, 0.74 and 1.21 cpd – adjusted *p* values = .006, <.001, and .002 respectively, Bonferroni corrected). Amplitudes were not significantly different at low corrugation frequencies and the highest corrugation frequency, showing that the difference was around the peak of the 1F1 tuning function. As expected, this shows that both the harmonic and the corrugation frequency determined the shape of the tuning functions.

The interaction between corrugation frequency and harmonic was also significant (*F*(3.66, 87.85) = 3.78, *p* = .009, *η^2^_G_* = 0.04 with Greenhouse-Geisser correction for sphericity as assessed by Mauchly’s test (*W* = 0.06, *p* < .001)). This implies that the shapes of the functions differed by harmonic, where the 1F1 response showed a high degree of tuning for corrugation frequency but the tuning was weak or absent in 1F2. Thus, 1F1 and 2F1 mechanisms were dissociated in a similar manner across both the RC1 and RC2 scalp components.

## DISCUSSION

Human disparity processing is subserved by multiple mechanisms differing in their corrugation frequency tuning, response dynamics, and cortical distribution. Harmonic analysis of the SSVEP grants access to underlying temporal channel dynamics, without depending on ‘transient’ or ‘sustained’ stimulus presentation profiles. Our results suggest links between sustained dynamics and relative disparity, and transient dynamics and absolute disparity, while providing a coarse functional localization of these signals.

### Relative disparity mechanisms drive the 1F1 response

Stereopsis is strongly dependent on the presence of references in the visual field and is optimal for a certain range of corrugation frequencies. We find that that the first harmonic of the changing disparity SSVEP shares these features, both at electrodes over early visual cortex (RC1) and electrodes over lateral occipital cortex (RC2). Adding binocular disparity information to a small fixation target amplified the first harmonic for both scalp components in our first experiment. This is reminiscent of the classic effects of adding references to stereograms, which results in an improvement in perceptual depth thresholds (Westheimer, 1979; McKee et al., 1990; Kumar and Glaser, 1991; Andrews et al., 2001).

The first harmonic response of RC1 is also strongly tuned for corrugation frequency, as is psychophysical disparity sensitivity across the wide range of studies shown in Figure 5, panel D. Disparity thresholds measured here are in the hyper-acuity range (Westheimer, 1979), being well under 1 arcmin under optimal conditions. Corrugation frequency tuning was also measured in the 1F1 response in RC2. Thus, the 1F1 signal is strongly driven by relative disparity mechanisms across multiple visual areas.

### Absolute disparity mechanisms and the 2F1 response

The evoked response also contains activity at the second harmonic. This component indexes transient responses that are equal across flat/disparate and disparate/flat stimulus transitions. The dependence of the second harmonic on references and corrugation frequency is less prominent than the first harmonic, especially for the RC1 component where its amplitude was unaffected by manipulating a binocular reference at fixation and its corrugation tuning was flat. Thus, we suggest that the 2F1 response is driven by mean changes in absolute disparity.

The 2F1 response amplitude was also largest for the full-field, absolute disparity condition in the corrugation frequency tuning experiment. Here, the DRDS alternated between disparate and non-disparate states, while in the grating condition the average disparity over the field was only one-half that. The boost in amplitude suggests that 2F1 is being driven by the local mean disparity, as would be the case if it were dominated by a contribution from cells’ responses to absolute disparity.

The second harmonic is unmeasurable in the RC2 component when only absolute disparity is available in the display. Thus, absolute disparity signals do *not* appear to be widely distributed. However, our stereograms were comprised of relatively small dots and small disparities. Studies using larger-field, coarse, absolute disparity stimuli have found early latency, disparity-tuned responses in macaque MT/MST (Takemura et al., 2000; Takemura et al., 2001; Takemura et al., 2002). The relative weighting of first and second harmonic responses in our system could be shifted by varying scale of the monocular half-images, or the disparity amplitude. Alternatively, signals may be invisible at the scalp if they are generated in cortical areas presenting a closed field configuration or are distant from the electrodes.

### Sustained and transient mechanisms in stereopsis

The relationship between sustained mechanisms and relative disparity has repeatedly been shown in psychophysical data, where longer stimulus durations result in lower depth detection thresholds (Ogle and Weil, 1958; Harwerth and Rawlings, 1977; Westheimer and Pettet, 1990). Manipulations of the *contrast* corrugation frequency content of disparity stimuli have linked relative disparity processing with sustained activity in the parvocellular pathway (Schor et al., 1984; Edwards et al., 1998; Kontsevich and Tyler, 2000; Gheorghiu and Erkelens, 2005; Lee et al., 2007).

Transient disparity mechanisms have been implicated by studies of the of vergence eye movements (Mitchell, 1970; Jones, 1980) where non-fusable disparate target can initiate brief, directionally appropriate vergence (Jones, 1980; Erkelens and Collewijn, 1985b, a). Consistent with this, depth sensations can be conveyed by very brief presentations of anticorrelated stimuli (Edwards et al., 1998; Schor et al., 1998; Pope et al., 1999), implicating a transient disparity system using correlation-based computations that do not depend on binocular feature matching (Doi et al., 2013).

The temporal dynamics of disparity tuned cells in V1 of macaque, by contrast, have been modelled by a single channel model comprising a bandpass linear monocular temporal kernel, followed by a rectifying non-linear binocular energy computation (Nienborg et al., 2005). This energy computation renders the temporal kernel of the disparity response monophasic and thus temporally low-pass, with the implication being that the response to disparity should have a lower high temporal frequency cut-off than the monocular kernel. This was observed in both single cells and their human psychophysical observers (see also (Richards, 1972; Regan and Beverley, 1973; Beverley and Regan, 1974; Norcia and Tyler, 1984; Gray and Regan, 1996).

A second implication of the Nienborg et al. model is that sensitivity to low temporal frequency disparity modulations should not fall off relative to moderate temporal frequencies. However, in three of four of their human observers, sensitivity was lower at 0.5 Hz than at 1-1.5 Hz, consistent with previous studies (Tyler, 1971; Richards, 1972; Gray and Regan, 1996; Lages et al., 2003). Nienborg et al.’s cellular measurements did not extend to 0.5 Hz, so the model prediction was not fully tested on the low-frequency range. Their model begs the question where the transient disparity responses measured in the oculomotor and behavioural literatures, and observed in own our VEP data, are purported to arise.

In our system, these predictions could be tested by measuring the temporal frequency tuning of the first and second harmonic response components. If the first harmonic reflects sustained processes, it should exhibit little low temporal frequency roll-off. Conversely, if the second harmonic response reflects a transient mechanism, its temporal tuning should be bandpass. The presence or absence of disparity references should also determine which component dominates.

### Purpose of duplex coding strategies in the disparity domain

Integration over long periods of time, say to increase acuity by averaging over noisy inputs, precludes detection of rapid changes that occur over shorter timescales. Sensory systems may solve this resolution/integration paradox through duplex coding systems that involve both transient and sustained mechanisms (Ikeda and Wright, 1972; Abraira and Ginty, 2013; Shiramatsu et al., 2016). In the visual system, transient and sustained channels originate in outputs of retinal ganglion cells (Cleland et al., 1971; Fukuda, 1971; Ikeda and Wright, 1972) that are segregated in the LGN and the input layers of V1, but converge thereafter (Nassi and Callaway, 2009).

These channels have been associated with distinct functional roles. As part of the magnocellular pathway, cells with transient response profiles support fixation and orientation behaviours, whilst cells with sustained response profiles in the parvocellular pathway support accurate registration of corrugation characteristics of the stimulus (Van Essen and Gallant, 1994; Kaplan and Benardete, 2001; Lee, 2011). In our experiments, the outputs of these channels do not vary over the stimulus conditions we employ because all of our monocular half-images are of identical spatiotemporal content. Despite this, we observe frequency domain responses originating purely from binocular signals that are consistent with transient and sustained temporal integration. Thus, a duplex coding strategy is recapitulated in the disparity domain itself, rather than being simply inherited from the monocular inputs.

### Topography of disparity responses

The approximate localization of the disparity signals we measure can be inferred from the topographies provided by Reliable Components Analysis. The midline occipital RC1 source is likely driven by signals in early visual areas including V1, V2 and V3, which contain cells tuned for absolute and/or relative disparity (Cumming and Parker, 1999; Thomas et al., 2002; Anzai et al., 2011). The proximity of V3B and its substantive contribution to form-from-disparity mechanisms (Kohler et al., 2019) is also likely to contribute to RC1.

The source of RC2 may lie in the LO complex where responses to dynamic changes in relative disparity have been measured using fMRI (Kohler et al., 2019). Although a form of relative disparity sensitivity has been demonstrated in macaque MT and MST (Krug and Parker, 2011), in human fMRI hMT+ is more commonly associated with absolute disparity processing (Neri et al., 2004; Neri, 2005; Orban et al., 2006; Parker, 2007) and is thus less likely to underlie RC2 activity.

RC2 is right lateralized here and in previous work using similar stimuli (Norcia et al., 2017). Consistent with this, depth discrimination is better in the left visual field (Durnford and Kimura, 1971; Grabowska, 1983) and several fMRI studies report right-lateralized disparity-related responses (Hirsch et al., 1995; Kwee et al., 1999; Nishida et al., 2001; Taira et al., 2001), although there is considerable inter-subject variation (Baecke et al., 2009).

### Conclusions

Our data suggest a duplex coding strategy for disparity in which a sustained channel processes the corrugation structure of the depth map, coupled with a transient channel in early visual cortex processing local disparity.

## Acknowledgements

This research was supported by grant no. EY018875 from the National Eye Institute, National Institutes of Health. The authors would like to thank Vladimir Vildavski and Alexandra Yakovleva for the development of instrumentation used in the experiments.

## References

Abraira VE, Ginty DD (2013) The sensory neurons of touch. Neuron 79:618–639.

Andrews TJ, Glennerster A, Parker AJ (2001) Stereoacuity thresholds in the presenc eof a reference surface. Vision Research 41:3051–3061.

Anzai A, Chowdhury SA, DeAngelis GC (2011) Coding of stereoscopic depth information in visual areas V3 and V3A. Journal of Neuroscience 31:10270–10282.

Baecke S, Lützkendorf R, Tempelmann C, Müller C, Adolf D, Scholz M, Bernarding J (2009) Event-related functional magnetic resonance imaging (efMRI) of depth-by-disparity perception: additional evidence for right-hemispheric lateralization. Experimental Brain Research 196:453–458.

Banks MS, Gepshtein S, Landy MS (2004) Why is spatial stereoresolution so low? Journal of Neuroscience 24:2077–2089.

Beverley KI, Regan D (1974) Temporal integration of disparity information in stereoscopic perception. Experimental Brain Research 19:228–232.

Bradshaw MF, Rogers BJ (1999) Sensitivity to horizontal and vertical corrugations defined by binocular disparity. Vision Research 39:3049–3056.

Bradshaw MF, Hibbard PB, Parton AD, Rose D, Langley K (2006) Surface orientation, modulation frequency and the detection and perception of depth defined by binocular disparity and motion parallax. Vision Research 46:2636–2644.

Brainard DH (1997) The Psychophysics Toolbox. Spatial Vision 10:433–436.

Burt P, Julesz B (1980) A disparity gradient limit for binocular fusion. Science 208:615–617.

Campbell FW, Maffei L (1970) Electrophysiological evidence for the existence of orientation and size detectors in the human visual system. Journal of Physiology 207:635–652.

Cleland BG, Dubin MW, Levick WR (1971) Sustained and transient neurones in the cat’s retina and lateral geniculate nucleus. Journal of Physiology 217:473–496.

Cottereau BR, McKee SP, Norcia AM (2012a) Bridging the gap: global disparity processing in the human visual cortex. Journal of Neurophysiology 107:2421–2429.

Cottereau BR, McKee SP, Ales JM, Norcia AM (2011) Disparity-tuned population responses from human visaul cortex. Journal of Neuroscience 31:954–965.

Cottereau BR, McKee SP, Ales JM, Norcia AM (2012b) Disparity-specific spatial interactions: evidence from EEG source imaging. Journal of Neuroscience 32:826–840.

Cumming BG, Parker AJ (1999) Binocular neurons in V1 of awake monkeys are selective for absolute, not relative, disparity. Journal of Neuroscience 19:5602–5618.

Didyk P, Ritschel T, Eisemann E, Myszkowski K, Seidel HP (2011) A perceptual model for disparity. ACM Transactions on Graphics (TOG) 30:1–10.

Dmochowski JP, Greaves AS, Norcia AM (2015) Maximally reliable spatial filtering of steady state visual evoked potentials. NeuroImage 109:63–72.

Doi T, Takano M, Fujita I (2013) Temporal channels and disparity representations in stereoscopic depth perception. Journal of Vision 13:1–25.

Durnford M, Kimura D (1971) Right hemisphere specialization for depth perception reflected in visual field differences. Nature 231:394–395.

Edwards M, Pope DR, Schor CM (1998) Luminance, contrast and spatial-frequency tuning of the transient-vergence system. Vision Research 38:705–717.

Erkelens CJ, Collewijn H (1985a) Motion perception during dichoptic viewing of moving random-dot stereograms. Vision Research 25:583–588.

Erkelens CJ, Collewijn H (1985b) Eye movements and stereopsis during dichopic viewing of moving random-dot stereograms. Vision Research 25:1689–1700.

Filippini HR, Banks MS (2009) Limits of stereopsis explained by local cross-correlation. Journal of Vision 9:8 1–18.

Fukuda Y (1971) Receptive field organization of cat optic nerve fibers with special reference to conduction velocity. Vision Research 11:209–226.

Gheorghiu E, Erkelens CJ (2005) Temporal properties of disparity processing revealed by dynamic random-dot stereograms. Perception 34:1205–1219.

Grabowska A (1983) Lateral differences in the detection of stereoscopic depth. Neuropsychologia 21:249–257.

Gray R, Regan D (1996) Cyclopean motion perception produced by oscillations of size, disparity and location. Vision Research 36:655–665.

Harwerth RS, Rawlings SC (1977) Viewing time and stereoscopic threshold with random-dot stereograms. American Journal of Optometry and Physiological Optics 54:452–457.

Haufe S, Meinecke F, Görgen K, Dähne S, Haynes JD, Blankertz B, Bießmann F (2014) On the interpretation of weight vectors of linear models in multivariate neuroimaging. NeuroImage 87:96–110.

Hess RF, Wilcox LM (1994) Linear and non-linear filtering in stereopsis. Vision Research 34:2431–2438.

Hess RF, Kingdom FAA, Ziegler LR (1999) On the relationship between the spatial channels for luminance and disparity processing. Vision Research 39:559–568.

Hirsch J, DeLaPaz RL, N R Relkin, Victor J, Kim K, Li T, Borden P, Rubin N, Shapley R (1995) Illusory contours activate specific regions in human visual cortex: evidence from functional magnetic resonance imaging. Proceedings of the National Academy of Sciences of the United States of America 92:6469–6473.

Hogervorst MA, Bradshaw MF, Eagle RA (2000) Spatial frequency tuning for 3-D corrugations from motion parallax. Vision Research 40:2149–2158.

Hou C, Gilmore RO, Pettet MW, Norcia AM (2009) Spatio-temporal tuning of coherent motion evoked responses in 4–6 month old infants and adults. Vision Research 49:2509–2517.

Ikeda H, Wright MJ (1972) Receptive field organization of ‘sustained’and ‘transient’ retinal ganglion cells which subserve different functional roles. Journal of Physiology 227:769–800.

Jones R (1980) Fusional vergence: Sustained and transient components. American Journal of Optometry 57:640–644.

Julesz B (1971) Foundations of Cyclopean Perception. Chicago: University of Chicago Press.

Kane D, Guan P, Banks MS (2014) The limits of human stereopsis in space and time. Journal of Neuroscience 34:1397–1408.

Kaplan E, Benardete E (2001) The dynamics of primate retinal ganglion cells. Progress in Brain Research 134:17–34.

Kleiner M, Brainard DH, Pelli DG (2007) What’s new in Psychtoolbox-3? Perception 36.

Kohler PJ, Cottereau BR, Norcia AM (2019) Image segmentation based on relative motion and relative disparity cues in topographically organized areas of human visual cortex. Scientifc Reports 9:1–18.

Kontsevich LL, Tyler CW (2000) Relative contributions of sustained and transient pathways to human stereoprocessing. Vision Research 40:3245–3255.

Krug K, Parker AJ (2011) Neurons in dorsal visual area V5/MT signal relative disparity. Journal of Neuroscience 31:17892–17904.

Kumar T, Glaser DA (1991) Influence of remote objects on local depth perception. Vision Research 31:1687–1699.

Kwee IL, Fujii Y, Matsuzawa H, Nakada T (1999) Perceptual processing of stereopsis in humans: High-field (3.0-tesla) functional MRI study. Neurology 53:1599–1601.

Lages M, Mamassian P, Graf EW (2003) Spatial and temporal tuning of motion in depth. Vision Research 43:2861–2873.

Lankheet MJ, Lennie P (1996) Spatio-temporal requirements for binocular correlation in stereopsis. Vision Research 36:527–538.

Lee B, Rogers B (1997) Disparity modulation sensitivity for narrow-band-filtered stereograms. Vision Research 37:1769–1777.

Lee BB (2011) Visual pathways and psychophysical channels in primate. The Journal of Physiology 589:41–47.

Lee S, Shioiri S, Yaguchi H (2007) Stereo channels with different temporal frequency tunings. Vision Research 47:289–297.

McKee SP, Levi DM, Browne SF (1990) The imprecision of stereopsis. Vision Research 30:1763–1779.

McKeefry DJ, Russell MHA, Murray IJ, Kulikowski JJ (1996) Amplitude and phase variations of harmonic components in human achromatic and chromatic visual evoked potentials. Visual Neuroscience 13:639–653.

Mitchell DE (1970) Properties of stimuli eliciting vergence eye movements and stereopsis. Vision Research 10:145–162.

Nassi JJ, Callaway EM (2009) Parallel processing strategies of the primate visual system. Nature Reviews Neuroscience 10:360–372.

Neri P (2005) A Stereoscopic Look at Visual Cortex. Journal of Neurophysiology 93:1823–1826.

Neri P, Bridge H, Heeger DJ (2004) Stereoscopic processing of absolute and relative disparity in human visual cortex. Journal of Neurophysiology 92:1880–1891.

Nienborg H, Bridge H, Parker AJ, Cumming BG (2004) Receptive field size in V1 neurons limits acuity for perceiving disparity modulation. Journal of Neuroscience 24:2065–2076.

Nienborg H, Bridge H, Parker AJ, Cumming BG (2005) Neuronal computation of disparity in V1 limits temporal resolution for detecting disparity modulation. Journal of Neuroscience 25:10207–10219.

Nishida Y, Hayashi O, Iwami T, Kimura M, Kani K, Ito R, Shiino A, Suzuki M (2001) Stereopsis-processing regions in the human parieto-occipital cortex. NeuroReport 12:2259–2263.

Norcia AM, Tyler CW (1984) Temporal frequency limits for stereoscopic apparent motion processes. Vision Research 24:395–401.

Norcia AM, Clarke M, Tyler CW (1985a) Digital filtering and robust regression techniques for estimating seonsory thresholds from the evoked potential. IEEE Engineering in Medicine and Biology Magazine 4:26–32.

Norcia AM, Sutter EE, Tyler CW (1985b) Electrophysiological evidence for the existence of coarse and fine disparity mechanisms in human. Vision Research 25:1603–1611.

Norcia AM, Gerhard HM, Meredith WJ (2017) Development of relative disparity sensitivity in human visual cortex. Journal of Neuroscience 37:5608–5619.

Ogle KN, Weil MP (1958) Stereoscopic vision and the duration of the stimulus. AMA Archives of Ophthalmology 59:4–17.

Olejnik S, Algina J (2003) Generalized eta and omega squared statistics: measures of effect size for some common research designs. Psychological Methods 8:434–447.

Orban GA, Janssen P, Vogels R (2006) Extracting 3D structure from disparity. Trends in Neurosciences 29:466–473.

Parker AJ (2007) Binocular depth perception and the cerebral cortex. Nature Reviews Neuroscience 8:379–391.

Pei F, Baldassi S, Tsai JJ, Gerhard HE, Norcia AM (2017) Development of contrast normalization mechanisms during childhood and adolescence. Vision Research 133:12–20.

Pelli DG (1997) The VideoToolbox software for visual psychophysics: transforming numbers into movies. Spatial Vision 10:437–442.

Peterzell DH, Serrano-Pedraza I, Widdall M, Read JCA (2017) Thresholds for sine-wave corrugations defined by binocular disparity in random dot stereograms: Factor analysis of individual differences reveals two stereoscopic mechanisms tuned for spatial frequency. Vision Research 141:127–135.

Pope DR, Edwards M, Schor CS (1999) Extraction of depth from opposite-contrast stimuli: transient system can, sustained system can’t. Vision Research 39:4010–4017.

Pulliam K (1982) Spatial frequency analysis of three-dimensional vision. Visual Simulation and Image Realism II 303:71–77.

Regan D, Beverley KI (1973) Some dynamic features of depth perception. Vision Research 13:2369–2379.

Regan D, Erkelens CJ, Collewijn H (1986) Necessary conditions for the perception of motion in depth. Investigative Ophthalmology and Visual Science 27:584–597.

Richards W (1972) Response functions for sine-and square-wave modulations of disparity. Journal of the Optical Society of America 62:907–911.

Rogers B, Graham M (1982) Similarities between motion parallax and stereopsis in human depth perception. Vision Research 22:261–270.

Schor CM, Wood IC, Ogawa J (1984) Spatial tuning of static and dynamic local stereopsis. Vision Research 24:573–578.

Schor CM, Edwards M, Pope DR (1998) Spatial-frequency and contrast tuning of the transient-stereopsis system. Vision Research 38:3057–3068.

Schumer R, Ganz L (1979) Independent stereoscopic channels for different extents of spatial pooling. Vision Research 19:1303–1314.

Serrano-Pedraza I, Read JCA (2010) Multiple channels ofr horizontal, but only one for vetical corrugations? A new look at the stereo anisotropy. Journal of Vision 10:1–11.

Shiramatsu TI, Noda T, Akutsu K, Takahashi H (2016) Tonotopic and field-specific representation of long-lasting sustained activity in rat auditory cortex. Frontiers in Neural Circuits 10:1–15.

Taira M, Nose I, Inoue K, Tsutsui K (2001) Cortical areas related to attention to 3D surface structures based on shading: An fMRI study. NeuroImage 14:959–966.

Takemura A, Inoue Y, Kawano K (2000) The effect of disparity on the very earliest ocular following responses and the initial neuronal activity in monkey cortical area MST. Neuroscience Research 38:93–102.

Takemura A, Inoue Y, Kawano K (2002) Visually driven eye movements elicited at ultra-short latency are severely impaired by MST lesions. Annals of the New York Academy of Sciences, 956:456–459.

Takemura A, Inoue Y, Kawano K, Quaia C, Miles FA (2001) Single-unit activity in cortical area MST associated with disparity-vergence eye movements: evidence for population coding. Journal of Neurophysiology 85:2245–2266.

Tang Y, Norcia AM (1995) An adaptive filer for steady-state evoked responses. Electroencephalography and Clinical Neurophysiology 96:268–277.

Thomas OM, Cumming BG, Parker AJ (2002) A specialization for relative disparity in V2. Nature Neuroscience 5:472–478.

Tyler CW (1971) Stereoscopic depth movement: Two eyes less sensitive than one. Science 174:958–961.

Tyler CW (1973) Stereoscopic vision: Cortical limitations and a disparity scaling effect. Science 181:276–278.

Tyler CW, Kontsevich LL (2001) Stereoprocessing of cyclopean depth images: horizontally elongated summation fields. Vision Research 41:2235–2243.

Van Essen DC, Gallant JL (1994) Neural mechanisms of form and motion processing in the primate visual system. Neuron 13:1–10.

Victor JD, Mast J (1991) A new statistic for steady-state evoked potentials. Electroencephalography and Clinical Neurophysiology 78:378–388.

Westheimer G (1979) Cooperative neural processes involved in stereoscopic acuity. Experimental Brain Research:585–597.

Westheimer G, Mitchell DE (1969) The sensory stimulus for disjunctive eye movements. Vision Research 9:749–755.

Westheimer G, Pettet MW (1990) Contrast and duration of exposure differentially affect vernier and stereoscopic acuity. Proceedings of the Royal Society of London Series B: Biological Sciences 241:42–46.

Wilcox LM, Allison RS (2009) Coarse-fine dichotomies in human stereopsis. Vision Research 49:2653–2665.

